# Implicating candidate genes at GWAS signals by leveraging topologically associating domains

**DOI:** 10.1101/087718

**Authors:** Gregory P. Way, Daniel W. Youngstrom, Kurt D. Hankenson, Casey S. Greene, Struan F. A. Grant

## Abstract

Genome wide association studies (GWAS) have contributed significantly to the understanding of complex disease genetics. However, GWAS only report associated signals and do not necessarily identify culprit genes. As most signals occur in non-coding regions of the genome, it is often challenging to assign genomic variants to the underlying causal mechanism(s). Topologically associating domains (TADs) are primarily cell-type independent genomic regions that define interactome boundaries and can aid in the designation of limits within which an association most likely impacts gene function. We describe and validate a computational method that uses the genic content of TADs to discover candidate genes. Our method, called “TAD_Pathways,” performs a Gene Ontology (GO) analysis over genes that reside within TAD boundaries corresponding to GWAS signals for a given trait or disease. We applied our pipeline to the GWAS catalog entries associated with bone mineral density (BMD), identifying ‘Skeletal System Development’ (Benjamini-Hochberg adjusted *p=*1.02x10^−5^) as the top ranked pathway. In many cases, our method implicated a gene other than the nearest gene. Our molecular experiments describe a novel example: *ACP2*, implicated at the canonical ‘*ARHGAP1*’ locus. We found *ACP2* to be an important regulator of osteoblast metabolism, whereas *ARHGAP1* was not supported. Our results via the example of BMD demonstrate how basic principles of three-dimensional genome organization can define biologically informed association windows.

## Introduction

GWAS have been applied to over 300 different traits, leading to the discovery of important disease associations^1^. However, assigning signals to causal genes has proven difficult because these signals fall principally within noncoding regions and do not necessarily implicate the nearest gene^2,3^. For example, a signal found in an *FTO* intron has been shown to physically interact with and lead to the differential expression of other genes, and not *FTO* itself ^4^. Moreover, there is evidence suggesting a type 2 diabetes GWAS association impacting *TCF7L2* also influences *ACSL5*^5^. It remains unclear how pervasive these kinds of associations are, but similar strategies are necessary in order for GWAS to better guide research and precision medicine^6^.

Chromatin interaction studies have discovered key genome organization principles including TADs^7^. TADs are genomic regions defined by increased contact frequency, consistency across cell types, and their enrichment of insulator element flanks^8^. Therefore, TADs can be used as boundaries of where non-coding causal variants will most likely impact tissue independent function.

We developed a computational approach, called “TAD_Pathways,” which uses TADs to determine candidate genes. We validated our method using BMD GWAS^9^. Using our method, we identified *ACP2* as a novel regulator of osteoblast metabolism.

## Methods

### Computational procedures to identify candidate genes

TAD_Pathways is a computational method that uses TADs to discover candidate genes (Figure 1A). Alternative approaches either assign genes based on nearest gene or by an arbitrary or a linkage disequilibrium-based window of several kilobases^10^ (Figure 1B). Here, we use human embryonic stem cell TAD boundaries as reported by Dixon et al. and converted to hg19 by Ho et al. to build TAD based genesets that consists of all Gencode genes that fall inside TADs implicated with BMD associations^8,11^. We perform a pathway overrepresentation test^12^ for the input TAD genes against Gene Ontology (GO) terms^13^. This determines if the geneset is associated with any term at a higher probability than by chance. We included both experimentally confirmed and computationally inferred genes, which permit the inclusion of putative genes which do not necessarily have literature support, but are predicted computationally. For validation, we consider only the most significantly enriched term, but a user can also select multiple. Our method also supports custom input SNP lists. TAD_Pathways software is available at https://github.com/greenelab/tad_pathways.

**Figure 1.**
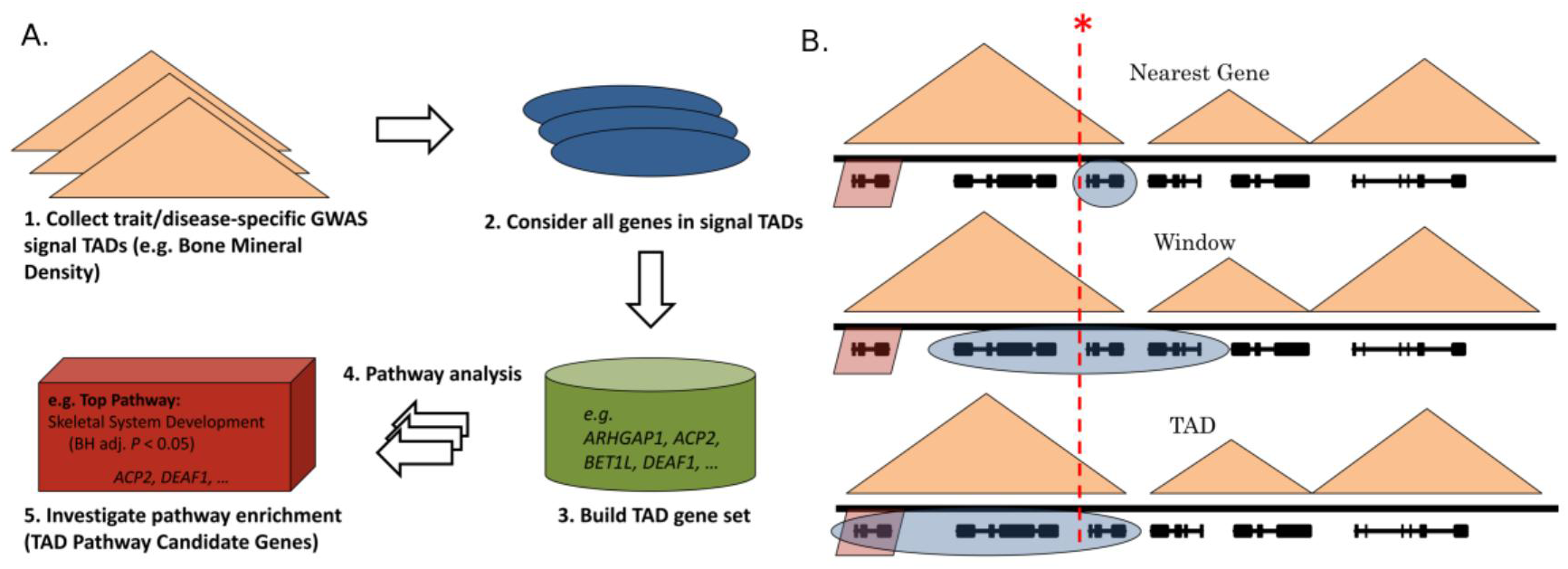
Concepts motivating our approach. Topologically associating domains (TADS)are shown as orange triangles, genes are shown as black lines, and a genome wide significant GWAS signal is shown as a dotted red line. **(A)** The TAD_Pathways method. An example using Bone Mineral Density GWAS signals is shown. **(B)** Three hypothetical examples illustrated by a cartoon. The ground truth causal gene is shaded in red. The method-specific selected genes are shaded in blue. The top panel describes a nearest gene approach. The nearest gene in this scenario is not the gene actually impacted by the GWAS SNP. The middle panel describes a window approach. Based either on linkage disequilibrium or an arbitrarily sized window, the scenario does not capture the true gene. The bottom panel describes the TAD_Pathways approach. In this scenario, the causal gene is selected for downstream assessment.

### Experimental knockdown of candidate genes

We investigated two candidate genes predicted by TAD_Pathways; *ACP2* and *DEAF1*. The corresponding BMD GWAS loci rs7932354 (cytoband: 11p11.2) and rs11602954 (cytoband: 11p15.5) are assigned to *ARHGAP1* and *BET1L,* respectively. The candidate genes were selected because no experimental support exists for their impact in bone and the nearest genes to the GWAS signals do not have immediate bone related functional implications. All experiments were conducted in three temporally separated independent technical replicates from cryopreserved P2 aliquots of a human fetal osteoblast cell line (ATCC hFOB 1.19 CRL-11372). Full experimental design can be found in the accompanying **Supplementary Information**.

siRNA transfections were conducted using a commercial reagent system with cells assigned to one of 8 experimental groups: untreated control, siRNA-negative transfection reagent control, scrambled control siRNA, *TNAP* siRNA, *ARHGAP1* siRNA, *ACP2* siRNA, *BET1L* siRNA or Suppressin (*DEAF1*) siRNA.

For quantitative gene expression analysis, whole RNA was harvested at Day 4, isolated and reverse-transcribed. Duplicate 20ng templates for each gene of interest were assayed via real-time PCR using SYBR reagents. Results were analyzed using the 2^-ddCt^ method using GAPDH as a housekeeping gene and reported as (mean±standard deviation) fold-change versus untreated controls. Cellular metabolism/proliferation was assessed at 1 and 4 days post-transfection using an MTT cell growth determination kit. Results are reported as the difference in mean absorbance at 570nm minus 690nm from Day 1 to Day 4 and error bars represent root-mean-square standard deviation from the measurements at both days. Early osteoblast differentiation was assessed at 4 days post-transfection using an alkaline phosphatase (ALP) kit. Dried plates were scanned and ALP+ intensities/areas were quantified in ImageJ. Values are reported as mean ± standard deviation.

## Results

### TAD_Pathways reveals candidate genes within phenotype-associated TADs

We applied TAD_Pathways to BMD GWAS results derived from replication-requiring journals^9,14–16^. Our method implicated ‘Skeletal System Development’ as the top ranked pathway (Benjamini-Hochberg adjusted *p*=1.02x10^−5^). For full BMD TAD_Pathways refer to **Supplementary Table S1**. Many candidates were not the nearest gene to the GWAS signal and several had independent eQTL support (**Supplementary Table S2**).

### siRNA Knockdown of TAD_Pathways Gene Predictions in Osteoblast Cells

We targeted the expression of four of these genes in vitro using siRNA and assessed transcriptional knockdown efficiency. Knockdown efficiencies were: TNAP 48.7±9.9% (*p*=0.141), *ARHGAP1* 68.7±14.3% (*p*=0.015), *ACP2* 48.9±6.4% (*p*=0.035), *BET1L* 56.4±1.0% (*p*>0.05) and *DEAF1* 52.7±9.2% (*p*=0.021) (Figure 2).

**Figure 2:**
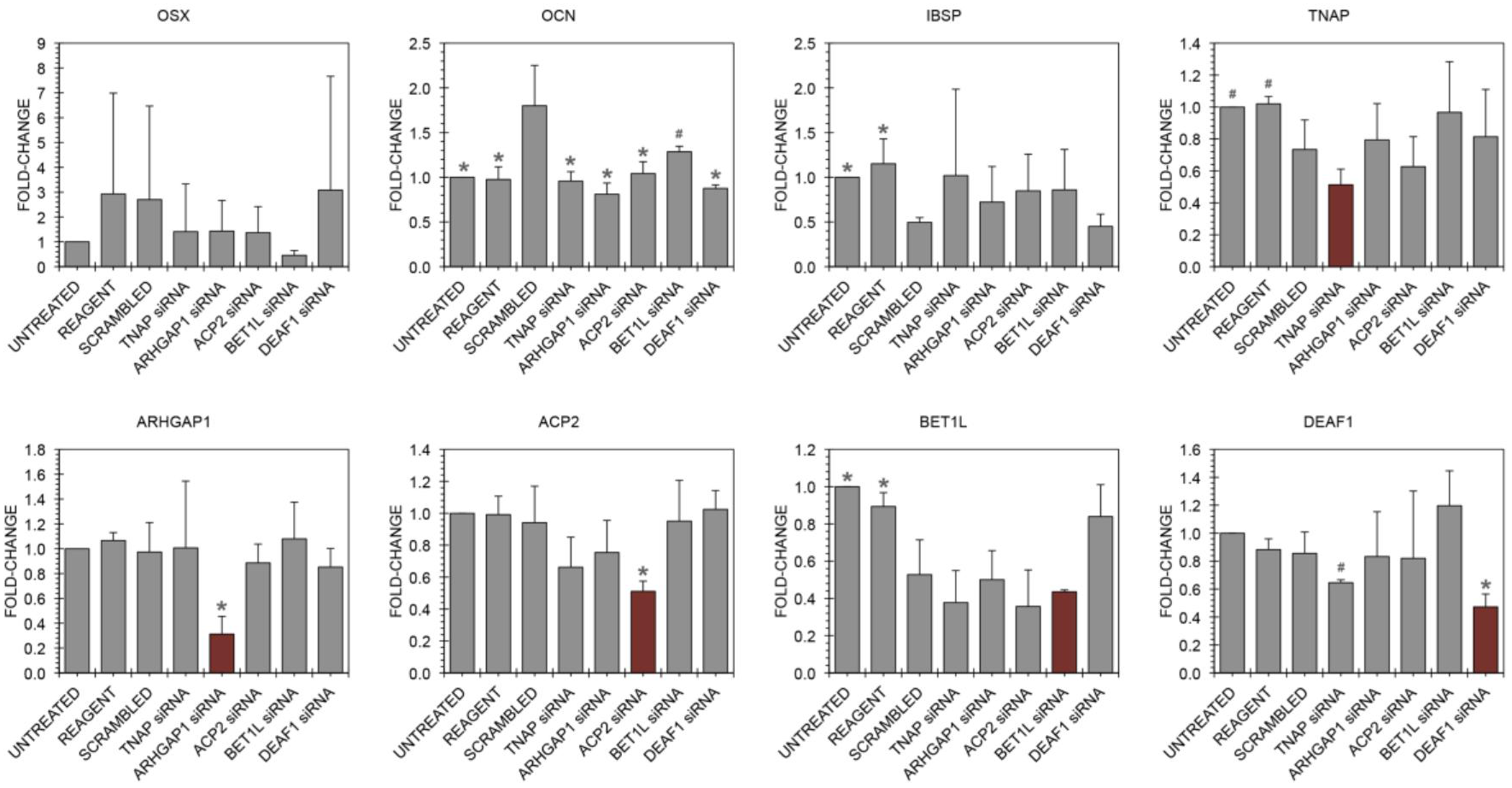
Real-time PCR of osteoblast differentiation genes and GWAS/TAD hits in hFOB cells. siRNA was used to knock down expression of *TNAP* (positive control), *ARHGAP1*, *ACP2*, *BET1L* and *DEAF1*. Relative expression of the osteoblast marker genes *OSX*, *OCN* and *IBSP* suggest that GWAS/TAD hits are not major regulators of bone differentiation in this model. Red bars highlight specificity of each siRNA knockdown. Values represent mean ± standard deviation. Statistical significance relative to the scrambled siRNA control is annotated as: \**p* ≤ 0.05 and #*p* ≤ 0.10 using a two-tailed Student’s *t*-test.

We noted variation across the three controls, with the scrambled siRNA control altering expression of *OCN* (osteocalcin), *IBSP* (bone sialoprotein), *TNAP* and *BET1L* (*p*<0.05). Relative to the scrambled siRNA control, *OCN* was downregulated in all siRNA groups (*p*<0.05) except for *BET1L* siRNA (*p*=0.122). *OSX*, *IBSP* and *TNAP* were not significantly altered by any siRNA treatment (Figure 2).

### Metabolic Activity of TAD_Pathways Gene Predictions

Treatment with *ACP2* siRNA led to a 66.0% reduction in MTT metabolic activity versus the scrambled siRNA control (*p*=0.012). *ARHGAP1* siRNA caused a 38.8% reduction, falling short of statistical significance (*p*=0.088). siRNA targeted against *TNAP*, *BET1L* or *DEAF1* did not alter MTT metabolic activity (Figure 3A).

**Figure 3.**
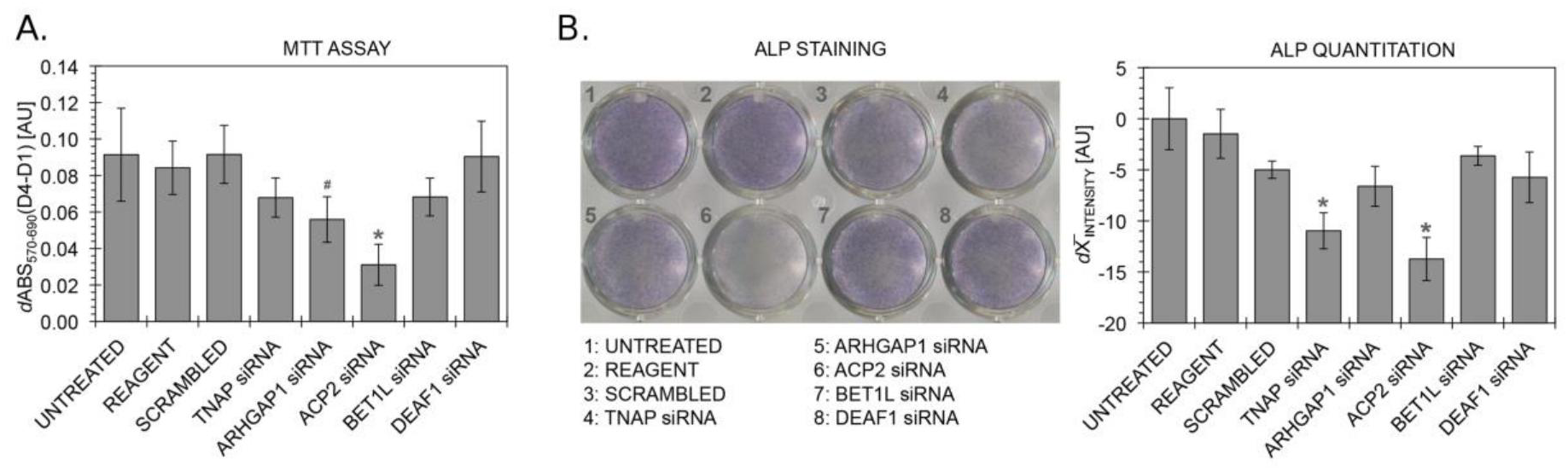
*Validating two ‘TAD Pathway’ predictions for Bone Mineral Density GWAS hits on hFOB cells.* siRNA was used to knock down expression of *TNAP, ARHGAP1, ACP2, BET1L* and *DEAF1*. **(A)** Knockdown of *ACP2* decreases cellular metabolic activity, demonstrated using an MTT assay. **(B)** ALP staining and quantitation indicates that knockdown of *TNAP* or *ACP2* inhibits performance in an osteoblast differentiation assay. Values represent mean ± standard deviation. Statistical significance relative to the scrambled siRNA control is annotated as: \**p* ≤ 0.05 and #*p* ≤ 0.10 using a two-tailed Student’s *t*-test.

### Alkaline Phosphatase Activity of TAD_Pathways Gene Predictions

ALP is highly expressed in osteoblasts; disruption of proliferation or osteoblast differentiation would result in downregulation of ALP. *TNAP* siRNA significantly reduced ALP intensity by 5.98±1.77 units versus the scrambled siRNA control (*p=*0.006). *ACP2* siRNA also significantly reduced ALP intensity by 8.74±2.11 (*p*=0.003). The control stained less intensely than untreated or transfection reagent controls, but this did not reach statistical significance (0.05<*p*<0.10) (Figure 3B).

## Discussion

We showed that TAD_Pathways can reveal functional gene to intermediate phenotype relationships using BMD. Several of the TAD_Pathways genes, such as *LRP5*, are *bona fide* BMD genes already identified by nearest gene GWAS, eQTL analyses, and clinical syndromes^17^, thus providing positive controls. However, several BMD GWAS signals do not have obvious nearest-gene associations with bone. Our results suggest that a nearby gene *ACP2,* and not the nearest gene *ARHGAP1*, regulates osteoblast proliferation/viability.

There are several limitations to the approach. Publication biases from pathway curation present challenges^18^. To lessen this bias, we include curated and computationally predicted GO annotations. Additionally, we used TAD boundaries defined by Dixon *et al.*, whereas increased Hi-C resolution reduced estimated TAD sizes^19^. Despite our method using larger TADs, thus including more presumably false positive genes, we still identify relevant pathways. Furthermore, the method will likely fail in a disease instigated by aberrant looping. We were also concerned that TAD_Pathways would work only in BMD. Indeed, when we applied TAD_Pathways to Type 2 Diabetes, we also identify several candidate genes that are not the nearest gene (see Supplementary Table S3). Moreover, the experimental validation was performed in a simplified *in vitro* cell culture system lacking organismal complexity, and the cell line selected is tetraploid, which may partially compensate for gene knockdown. While TAD_Pathways identified several candidate genes, we only examined two, and our validation approach does not directly interrogate each SNP. Lastly, one of the investigated GWAS SNPs, rs7932354, located in the *ARHGAP1* promoter, is an *ARHGAP1* eQTL in several tissues as determined by GTEx^20^. However, none of these tissues are bone related and our screen did not implicate *ARHGAP1* in osteoblast processes.

In conclusion, TAD_Pathways can be used as a candidate gene discovery tool that uses chromatin looping to map associations to candidate genes. We also validated our method by implicating *ACP2* as a gene involved in BMD determination, which warrants future investigation. We believe TAD_Pathways and algorithms that leverage 3D genomic structure will be poised to discover novel disease features.

## Acknowledgements

Hannah E. Sexton and Troy L. Mitchell assisted in optimizing siRNA transfection conditions. Daniel Himmelstein and Amy Campbell performed analytical code review.

## Availability of data and material

All data used to construct the TAD_Pathways approach are publically available datasets. We make all software used to develop this approach publically available in a GitHub repository (http://github.com/greenelab/tad_pathways). We also provide a docker image (https://hub.docker.com/r/gregway/tad_pathways/) and archive the GitHub software on Zenodo (https://zenodo.org/record/163950).

